# Application of RFdiffusion to predict interspecies protein-protein interactions between fungal pathogens and cereal crops

**DOI:** 10.1101/2024.09.17.613523

**Authors:** Olivia C. Haley, Stephen Harding, Taner Z. Sen, Margaret R. Woodhouse, Hye-Seon Kim, Carson Andorf

## Abstract

Plant pathogenic fungi secrete small proteins known as effectors which help overcome the plant defense response and cause disease. The concept of effector-triggered immunity in plants evolved from the “gene for gene hypothesis” which describes plant resistance or susceptibility to plant pathogens based on interspecies protein-protein interactions (PPIs) between plant-derived resistance (R) genes and pathogen-derived avirulence (Avr) effector genes. Understanding the molecular interactions mediating these host-pathogen interactions in effector-triggered immunity is thus essential to managing fungal disease. *In silico* methods of predicting interspecies PPIs have been heavily studied to identify target genes for crop resistance. But conventional sequence-based homology methods (i.e., interlog, domain-based inference) for predicting interspecies PPIs are not as powerful as methods that also incorporate structural homology. The objective of this study was to develop a computational workflow to predict PPIs between pathogenic fungi and their cereal hosts by leveraging recent advances in artificial intelligence and structural biology. This workflow proposes the use of a generative model, RFdiffusion, to predict the structure of truncated segments of proteins likely to bind to query effector proteins. The binder structures were filtered based on the number of contacts at the effectors’ known binding residues. Acceptable structures were then input into FoldSeek to search the host proteome for host proteins containing similar sub-structures. Experimentally-validated PPIs between rice (*Oryzae sativa* cv. ‘Japonica’) and rice blast fungus (*Magnaporthe oryzae*) were used for workflow validation. The effects of binder length and the binding residues’ mode of action (i.e., residues at active/substrate recognition sites) on the binder quality and presumptive host protein matches were explored. Ultimately, 11 out of 14 experimentally validated PPIs were recovered computationally, indicating a high recall (>78%) for the workflow. The shorter binders recovered most of the PPIs, but may have produced the most false positives, as functional analyses revealed that these host proteins displayed a wide variety of functions. These findings emphasize that subject matter expertise is still required to decipher the prediction results. Yet, this framework for elucidating interactions between fungal pathogens and host proteins could provide valuable insight into mechanisms of susceptibility or resistance at a scale friendly to limited computational resources, and facilitate the development of control strategies that reduce crop diseases.

## INTRODUCTION

Fungal plant pathogens are a major cause of widespread agricultural losses, particularly in economically important cereal crops like rice, wheat and maize (Liu et al., 2022). Beyond agricultural and economic losses, some of these pathogens pose serious food safety concerns because they can produce mycotoxins and mycotoxin derivatives in grains that are harmful to human and animal health when consumed (Zhang et al., 2020). With climate change creating more favorable conditions for fungal growth, the niche of mycotoxigenic species is likely to expand (Chhaya et al., 2022). Therefore, continued research and development of technologies that predict and detect potential threats from emerging fungal pathogens is critical for developing control strategies and mitigating crop loss caused by fungal disease.

Fortunately, there have been long-standing scientific efforts and strides made towards developing such technologies, especially through studying protein function. In the plant immune response, proteins play a critical role in pathogen perception, cell signaling cascades, and oxidative bursts (Nishad et al., 2020). Fungal proteins called effectors, are secreted by pathogens to manipulate host cell processes, facilitate host colonization, evade host defenses, and to produce mycotoxins (Liu et al., 2022).

Effector-triggered immunity evolved from the “gene for gene hypothesis” which describes the interaction between plant-derived resistance (R) genes and pathogen derived avirulence (Avr) effector genes (Kumar et. al., 2022). The critical role of proteins in the plant immune response and fungal infection has led to their use as biomarkers for developing pesticides (Gressel, 2022), biosensors and immunoassays (Oliveira et al., 2019), designer plant proteins (Césari et al., 2022; Godana et al., 2023) and targeted crop breeding (Dracatos et al., 2023).

Understanding how fungal proteins affect plant metabolism is essential for maximizing the efficacy of protein-based and chemical agricultural innovations. Many of these proteins affect metabolism through intra- and interspecies protein-protein interactions (PPIs)(Liu et al., 2022). Thus, predicting this cross-talk has been a prime target for machine learning (Kaundal et al., 2022; Khairi et al., 2023; Lei et al., 2023; Yang et al., 2019; Zheng et al., 2021). *Intra*species PPI networks have been developed *in silico* for a variety of cereal crops (Ma et al., 2019; Musungu et al., 2015) and plant pathogens (Guo et al., 2013; Zhang et al., 2017), which have improved our understanding of fungal effectors (Yang et al., 2019) and reduced the time and cost needed to identify candidate genes for plant pathogen resistance (Zheng et al., 2021). However, predicting *inter*species PPIs between crop plants and fungal pathogens can be a much more arduous task, and falls far behind that in mammals and/or human research for various reasons.

Data availability is a major obstacle at the front-end of interspecies PPI data analyses. Most PPI databases (i.e., STRINGdb, MINT, IntAct) predominantly contain interactions from human or model animal systems (i.e., rat, Drosophila), and their pathogens of interest (Yang et al., 2019). Of the interspecies PPI data readily available in plants, the majority is from the model organism *Arabidopsis thaliana* (Yang et al., 2019). The number of experimentally resolved plant protein structures is also very low, with proteins from *Viridiplantae* (‘green plants’; Taxon ID: 33090) accounting for less than four percent of all available experimental structures in the Protein Data Bank (PDB) (Burley et al., 2023). In the past, sequence similarity-based methods such as interlog and domain-based inference were primarily used to infer PPIs by transferring assigned protein function from well-studied model organisms to the study’s target organism (Yang et. al, 2019). But using sequence similarity may only be effective for sequences that share a high degree of similarity (Mika & Rost, 2006). In fungi, recent work highlights several classes of effectors (i.e., MAX effectors, ToxA-like family) whose members share high structural similarity and phenotypic outcomes, but low sequence similarity (Jones et al., 2021). Once more, a study on the evolutionary dynamics of resistance (R) genes in plants show that the sequence-based homologs of R-genes, such as the paired R-genes Pik1/Pik2, may actually encode pseudogenes (Mizuno et al., 2020). Such findings may explain why the rice reference genome (cultivar Nipponbare) is not resistant to rice blast fungus despite having the Pik1/Pik2 homologs Pik5-NP/Pik6-NP.

However, there are also obstacles to using structural similarity or biochemistry alone for predicting interspecies PPIs. Even effectors with a high degree of structural similarity may have different surface charge distributions caused by amino acid substitutions, ultimately affecting the specificity of their interactions with plant host receptors (Zhang et al., 2018). For example, the effector AvrPib is structurally similar to MAX effectors (AvrPikD, AvrPia, etc.), except the protein has a distinct, positively-charged patch and hydrophobic core that impacts its avirulence function, nuclear localization, and likely how it is recognized (Zhang et al., 2018).Evaluating potential PPIs based on the structures’ binding affinity may also be less accurate since the binding affinity of interacting effector-host protein domains is not always strong, but may be moderate (i.e., AvrPia/RGA5; Ortiz et al., 2017) to weak (AvrPia/Pik1; Varden et al., 2019), especially for receptors with generalized recognition roles within the plant. Combining recent advances in both sequence and structure-based machine learning models could lead to more accurate interaction predictions and intriguing discoveries (Khairi et al., 2023; Yang et al., 2019; Zheng et al., 2021).

State-of-the-art computational methods like AlphaFold (Jumper et al., 2021), RoseTTA fold (Baek et al., 2021), and ESMFold (Lin et al., 2023) have opened the door for 3D protein structure-based inference, and their applicability to PPI prediction is highly promising. This study builds on this approach and explores the use of RFdiffusion, a generative model based on RoseTTA fold that has undergone fine-tuning for the generation of proteinaceous binders using only a query proteins’ structure (from experimentation or AlphaFold predictions) and the residues where the interaction is most likely to occur (‘hotspots’; Watson et al., 2023). The novelty of RFdiffusion is its use of denoising diffusion probabilistic models (DDPMs) which are trained to denoise data corrupted with Gaussian noise, and allows the workflow to generate realistic backbone structures from the extremely large protein space (Watson et al., 2023; Wu et al., 2022; Yim et al., 2023). Another major benefit of the RFdiffusion pipeline is that it also includes Protein-MPNN (Dauparas et al., 2022), a robust deep learning-based protein sequence design method to predict the amino acid sequence corresponding to the generated backbone, and AlphaFold to fold and validate the predicted binder structures. From the perspective of PPI prediction, such diffusion models could be used to produce a balance of existing and novel protein structures to not only study, but develop new protein-based and chemical technologies. However, the use of diffusion to predict PPIs has not yet been comprehensively studied in plants.

Despite its very strong results in protein binder design, RFdiffusion has been used in relatively few published studies at the time of this publication (Harris et al., 2024; Vázquez Torres et al., 2024; Liu et al., 2024). Thus, there is limited information regarding best practices to use RFdiffusion for the objective of predicting interspecies PPIs. Furthermore, use-cases in organisms that are under-represented in the protein sequence space (i.e., UniProtKB) and/or protein structure databases (i.e., PDB) are lacking. This study aims to apply RFdiffusion to generate proteinaceous binders for fungal effector proteins, hypothesizing that the sampling process will uncover native, interacting host plant proteins.

## MATERIALS AND METHODS

The steps in this approach are summarized in Figure 1.

**Figure 1.**
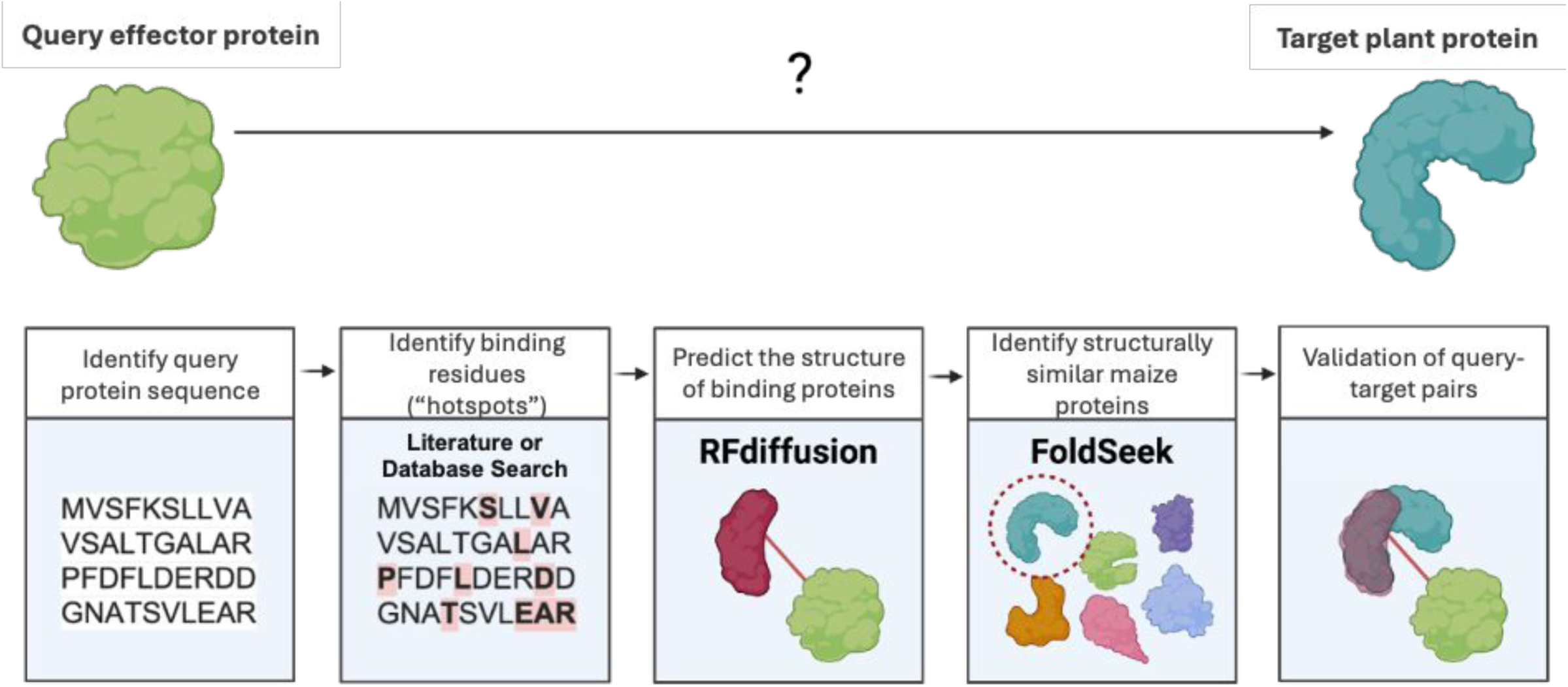
A summary of the workflow used in this study. Protein binding residues (hotspots) were extracted from the literature or from databases of conserved protein binding domains (i.e., InterPro). The experimentally validated or predicted hotspots were then passed to RFdiffusion to generate protein binders. Since RFdiffusion may not use all the hotspots during diffusion, the binders were filtered based on the number of interaction hotspots between the binder and query protein. The rice protein space was then searched for proteins with high structural similarity to the binders using FoldSeek. Illustration created using the Biorender platform.

### Validation dataset

Effectors from the rice blast fungus *Magnaporthe oryzae* were used to validate the workflow (*Table 1*). *M. oryzae* effectors were chosen for this study as they are relatively well-characterized, and the rice/rice blast pathosystem is well-studied. Thus the effectors’ interacting rice proteins and active sites were easily retrieved from the literature (see *Table 1*). To query the host proteome, a target database was compiled in FoldSeek using the predicted AlphaFold2 (AF2) structures of rice proteins downloaded from the AlphaFold Protein Structure Database (https://alphafold.ebi.ac.uk/; Varadi et al., 2024) using the reference proteome (UP000059680; *Oryza sativa* subsp. Japonica, cv. Nipponbare). Of note, the R-genes Pik1 and Pik2 are required for avirulence protein recognition in resistant cultivars like cv. Kusuabe. Within the nonresistant Nipponbare cultivar, the proteins corresponding to Pik5-NP and Pik6-NP were used as equivalents to represent the pair as per the findings of Zhai et al. (2010).

**Table 1.**
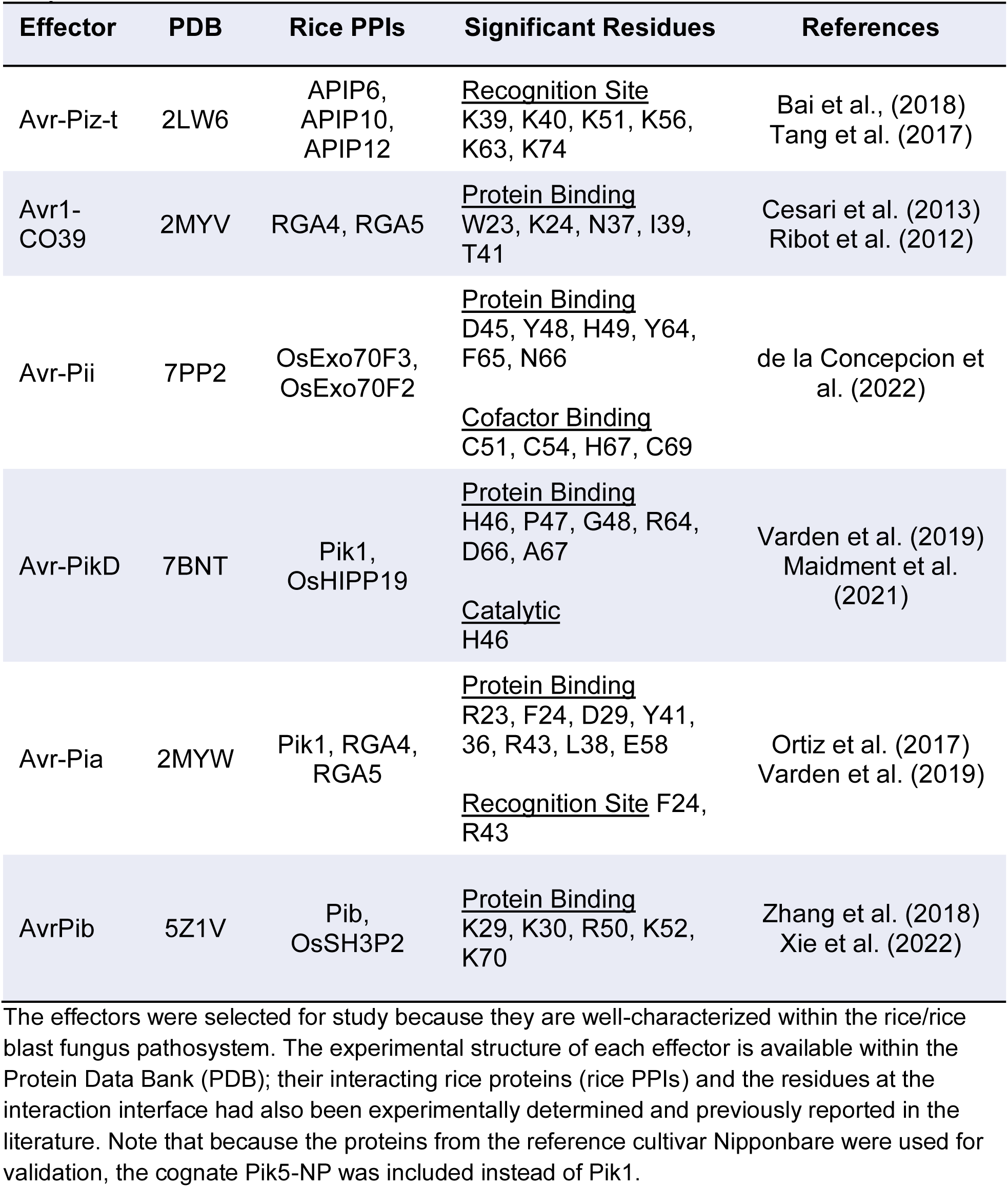
An overview of the *Magnaporthe oryzae* effectors used for workflow validation in this study.

### Binder Generation with Diffusion

The interacting residues and a PDB file for each effector were used as input for RFdiffusion (https://github.com/RosettaCommons/RFdiffusion) to predict the structure of proteinaceous binders. The beta model was used to improve the balance between alpha helix and beta sheet secondary structures seen in the generated binders, and the default number of iterations was used. The binder length and hotspot mode of action were explored for their effect on false positive/negative hits and the recall of the experimentally validated proteins in rice. The lengths of the binders (contigs) were either 75, 100, or 150 amino acids. These lengths were chosen based on data from the original publication; shorter binders generated during unconditional monomer design tended to have higher quality metrics (TM score, r.m.s.d). Other studies have also found success in their various objectives using short (typically ≤100 residues) binders (Liu et al., 2024; Vázquez Torres et al., 2024). Moreover, the results of this study will show that surpassing 150 amino acids in length results in a higher number of false negatives.

Based on the literature (*Table 2*), the hotspot mode of action was broadly categorized into protein-binding, cofactor binding, catalytic site, or recognition site. Protein-binding is the most general, including all the residues where an interaction occurs. Cofactor binding indicates that the residue participates in binding of a metallic or non-protein cofactor. Residues are considered a recognition site if there is evidence that it participates in binding to a receptor protein, leading to the propagation of the plant response. Catalytic residues directly participate in the catalysis of the ligand. For each contig length and mode of action, one hundred binders were generated, as this number is manageable for groups with limited computing resources. This process is facilitated by the availability of an RFdiffusion notebook on the Google Colab platform.

**Table 2.** A summary of the generated binders and recovered protein-protein interactions from the *M. oryzae* effectors using the workflow.

| Effector | PPIs | Site | 75 | 100 | 150 |
| --- | --- | --- | --- | --- | --- |
| Avr-Piz-t | APIP6, **APIP10**, APIP12 | Protein Binding | 2/62 | 0/56 | 0/57 |
| Avr1-CO39 | **RGA4, RGA5** | Protein Binding | 5/71 | 0/64 | 0/48 |
| Avr-Pii | **OsExo70F2, OsExo70F3** | Protein Binding | 2/95 | 5/93 | 6/90 |
|  |  | Cofactor Binding | 2/64 | 0/54 | 2/54 |
| Avr-PikD | **Pik1 (Pik5-NP), OsHIPP19** | Protein Binding | 8/91 | 3/81 | 0/69 |
|  |  | Catalytic Site | 3/76 | 1/71 | 0/70 |
| Avr-Pia | **Pik1 (Pik5-NP), RGA4, RGA5** | Protein Binding | 2/75 | 0/75 | 0/66 |
|  |  | Recognition Site | 1/60 | 1/72 | 0/61 |
| AvrPib | **Pib**, OsSH3P2 | Protein Binding | 0/90 | 1/91 | 0/77 |

The experimentally determined rice PPIs in bold are those that were detected (recovered) by the workflow, whereas un-bolded PPIs were not recovered. The fields 75, 100, and 150 refer to the amino-acid length of the binders. During the structural similarity search in FoldSeek, query binder – target host protein matches were only retained if the pair had a structural bit score $\geq 50$ . For each length and type of residues passed during diffusion (site), the numerator is the number of generated binders which returned an anticipated PPI; the denominator is the number of generated binders which were acceptable – thus passed to the structural similarity search.

RFdiffusion outputs a PDB file for each binder which shows the complex formed between the input query effector and the predicted binder. According to the RFdiffusion GitHub repository, ‘Of all of the hotspots which are identified on the target, 0-20% of these hotspots are actually provided to the model and the rest are masked.’ In short, RFdiffusion randomly chooses between 0 to 20% of the input hotspots to generate potential binders. Thus, for initial filtering, complexes with few (typically less than 2) to no interactions at the specified hotspots were excluded from further analysis. An interaction was deemed likely to occur if the distance between the binder’s alpha carbon (C_α_) and the query effectors’ beta carbons (C_β_) was less than 10 Å, based on previous studies on the most accurate residue distance metrics for detecting contacts (Bolser et al., 2008; Iyer et al., 2020) and data on the recall of PPIs according to different distance thresholds (*Table 3*). While the C_β_–C_β_ distance is the most accurate metric, the distance between binder C_α_ and query effectors C_β_ was used because RFdiffusion generates structures with all glycine residues. Thus, the binders may reflect a plausible secondary structure, but with inaccurate amino acid side chains.

**Table 3.** A summary of how the threshold on the distance between the effectors’ alpha carbon and binders’ beta carbon affected the number of acceptable binders and positive PPIs.

| Effector | Site | $\leq 8 \text{ \AA}$ | | | $\leq 10 \text{ \AA}$ | | | $\leq 12 \text{ \AA}$ | | | $\leq 14 \text{ \AA}$ | | |
| --- | --- | --- | --- | --- | --- | --- | --- | --- | --- | --- | --- | --- | --- |
|  |  | 75 | 100 | 150 | 75 | 100 | 150 | 75 | 100 | 150 | 75 | 100 | 150 |
| Avr-Piz-t | PB | 1/24 | 0/15 | 0/15 | 2/62 | 0/56 | 0/57 | 3/90 | 0/89 | 0/78 | 3/94 | 0/94 | 0/90 |
| Avr1-CO39 | PB | 5/68 | 0/58 | 0/40 | 5/71 | 0/64 | 0/48 | 5/73 | 0/71 | 0/58 | 5/84 | 0/82 | 0/74 |
| Avr-Pii | PB | 1/86 | 5/88 | 5/86 | 2/95 | 5/93 | 6/90 | 2/96 | 5/96 | 6/94 | 2/96 | 5/96 | 6/95 |
|  | CFB | 1/32 | 0/34 | 1/39 | 2/64 | 0/54 | 2/54 | 2/68 | 0/64 | 3/66 | 2/77 | 2/72 | 4/69 |
| Avr-PikD | PB | 7/84 | 3/74 | 0/65 | 8/91 | 3/81 | 0/69 | 8/91 | 3/81 | 0/69 | 8/91 | 3/81 | 0/70 |
|  | CAT | 3/49 | 1/48 | 0/56 | 3/76 | 1/71 | 0/70 | 3/77 | 1/71 | 0/70 | 3/77 | 1/71 | 0/71 |
| Avr-Pia | PB | 2/55 | 0/59 | 0/46 | 2/75 | 0/75 | 0/66 | 2/83 | 0/81 | 1/75 | 2/89 | 0/90 | 1/85 |
|  | REC | 0/32 | 0/36 | 0/26 | 1/60 | 1/72 | 0/61 | 1/61 | 1/77 | 0/65 | 2/67 | 1/78 | 1/71 |
| AvrPib | PB | 0/58 | 1/56 | 0/54 | 0/90 | 1/91 | 0/77 | 0/93 | 1/94 | 0/84 | 0/94 | 1/96 | 0/84 |

The fields 75, 100, and 150 refer to the length of the binders in amino acid residues. The sites are abbreviated; PB - protein binding, CFB - cofactor binding, CAT - catalytic site, REC - substrate recognition site. The fractions indicate the number of generated binders which returned the anticipated PPI match (numerator; bit score $\geq 50$ ) and the number of generated binders which were acceptable – thus passed to the structural similarity search – (denominator) is also shown for each site and binder length.

The structures of the acceptable binders were extracted from the PDB files using custom scripts in Biopython (ver. 1.83) and passed to FoldSeek to query the custom host databases and identify target proteins whose structure was similar to the binder. Once again, because RFdiffusion generates structures with all glycine residues, only parameters for comparing structural similarity were used in the FoldSeek easy-search function. The short length of the binders relative to potential matches also warranted certain parameters to reduce the number of false positives. Specifically, the alignment was performed based on a minimum 90% coverage of the query binder using the TMalign algorithm (Zhang & Skolnick, 2005) within FoldSeek (--alignment-type 1 --cov-mode 2 -c 0.90). Only proteins with a structural bit score ≥ 50 (the recommended threshold; Barrio-Hernandez et al., 2023) were included in subsequent analyses.

### Functional Enrichment Analysis

To understand the diversity of the presumptive PPIs during validation, gene ontology (GO) enrichments of the rice target proteins were performed using GOATOOLS (Klopfenstein et al., 2018) with annotations from the Gene Ontology Meta Annotator for Plants (GOMAP; Wimalanathan et al., 2021) as the ground truth. We applied the Bonferroni correction to account for multiple comparisons.

## RESULTS

### Overall workflow performance on the validation dataset

A summary of the workflow’s performance for the validation *M. oryzae* effectors is provided in *Table 2*. The workflow captured 11 out of the 14 experimentally-determined PPIs between the *M. oryzae* effectors and rice proteins. APIP6 and APIP12 were likely not recovered because RFdiffusion doesn’t generate backbones with extensive disordered regions, such as those in their AF2 structures that would have prevented high confidence detection during the structural similarity search.

### The impact of binder length and active site type on the workflow performance

Due to the limited information on best practices, the binder length, number of generated backbone structures, and active site type were explored during validation for their downstream impact on the structural similarity search and recall of protein targets (*Figure 2, 3*). We hypothesized that shorter binders may lead to high levels of false positives (‘noise’) since they increase the chance of matches occurring at a structure that is shared between a wide variety of proteins. Meanwhile longer binders may incorporate multiple structural domains that are possible, but ultimately exclude protein families important to the study’s objectives (increase false negative rate).

**Figure 2.**
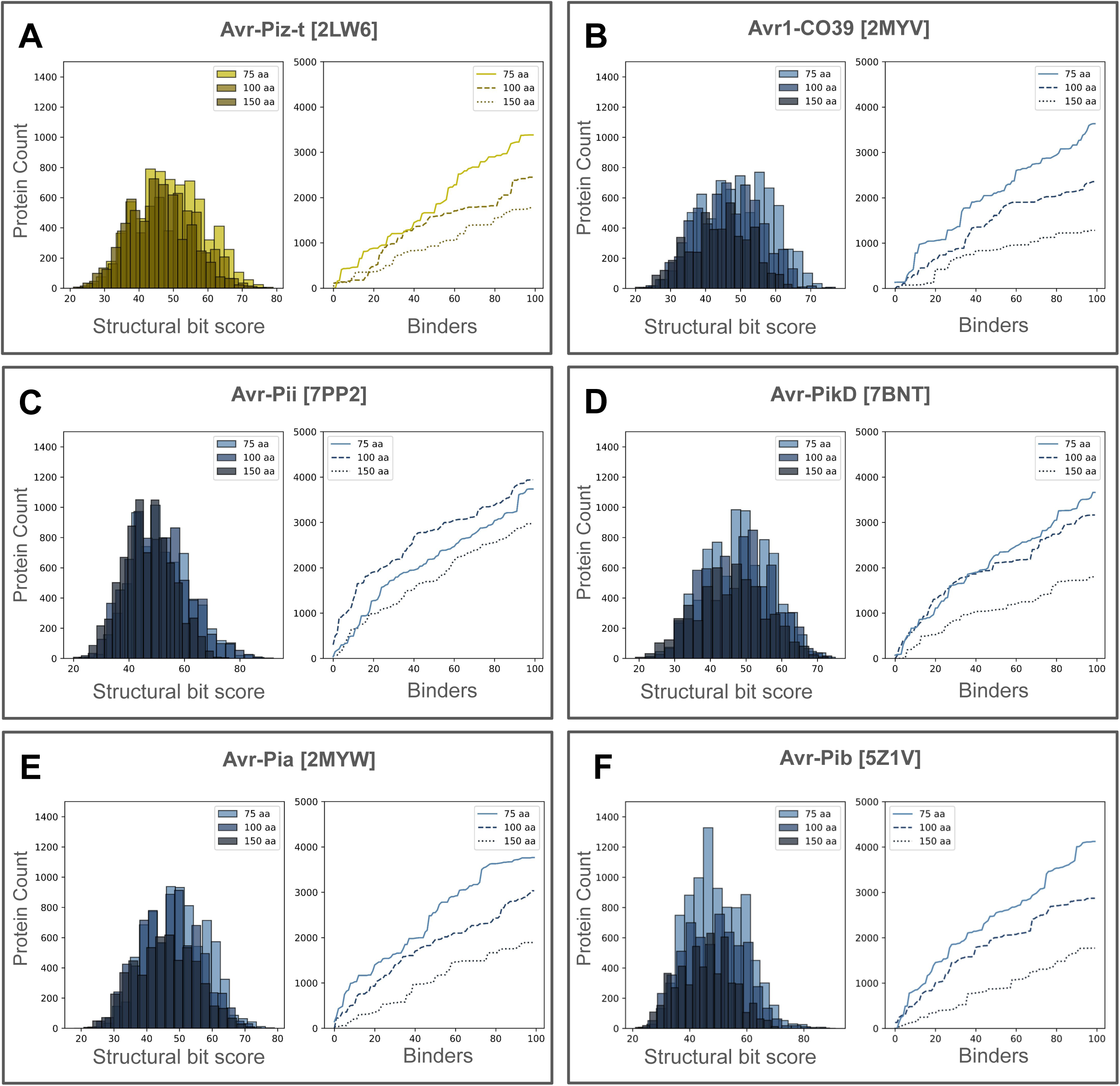
The results for *M. oryzae* validation proteins according to the protein binding residues that were used as input for RFdiffusion. Each panel shows the overall distribution of TM-scores obtained from comparing the binders against a database of proteins from *Oryzae sativa* in FoldSeek, by the amino acid length of the binder (left). The number of unique proteins in response to the increasing number of binders is shown (right) for the 75, 100, and 150 amino acid long binders.

**Figure 3.**
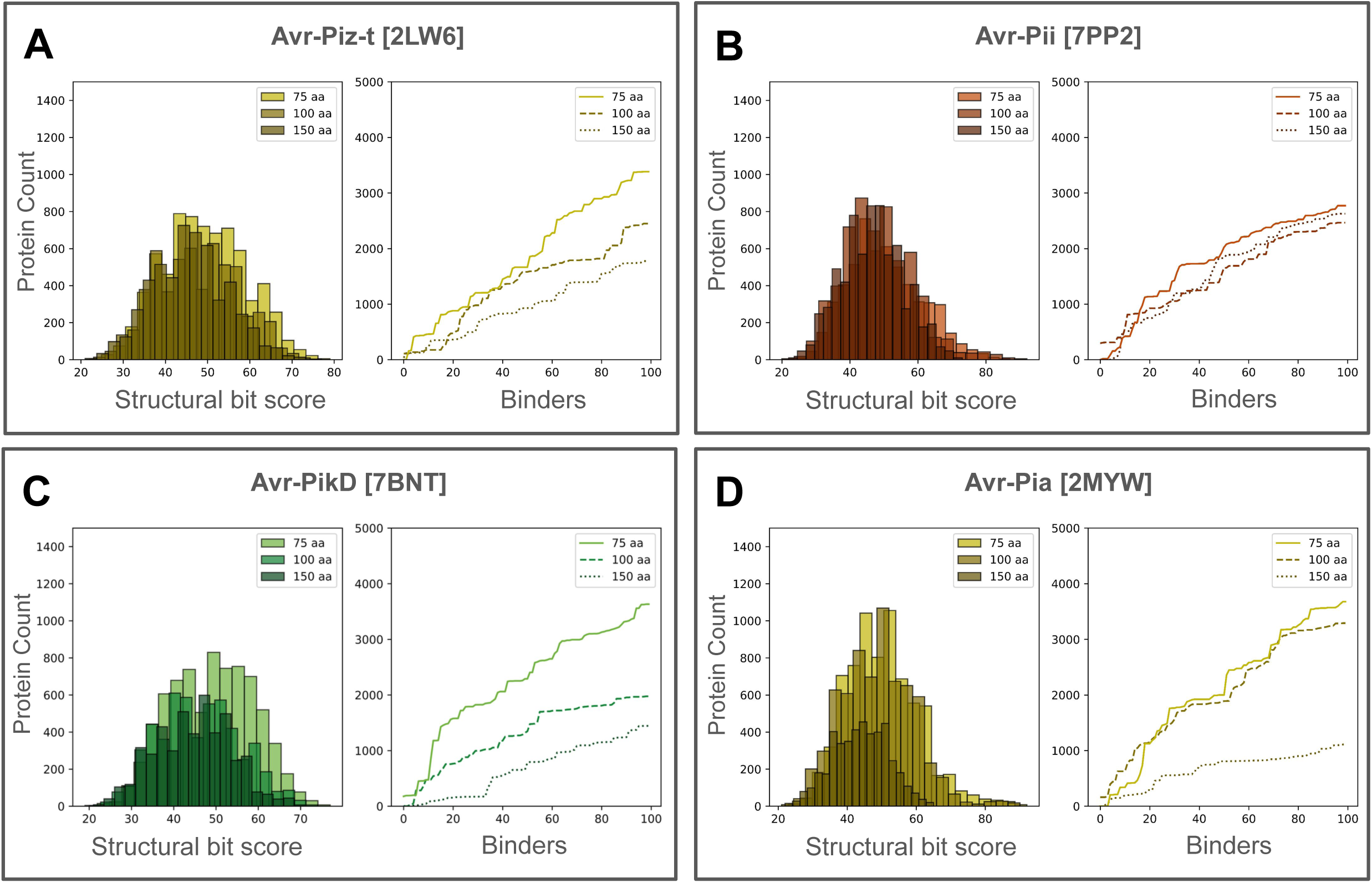
The results for *M. oryzae* validation proteins according to the protein binding residues that were used as input for RFdiffusion. Each panel shows the overall distribution of TM-scores obtained from comparing the binders against a database of proteins from *Oryzae sativa* in FoldSeek, by the amino acid length of the binder (left). The number of unique proteins in response to the increasing number of binders is shown (right) for the 75, 100, and 150 amino acid long binders.

Indeed, the 75 amino acid (aa)-long binders tended to confer the greatest number of structurally similar protein matches that also met the threshold (bit score ≥ 50) followed by the 100 and 150 aa-long binders, respectively (*Figure 2, 3*); AVR-Pii was the exception (*Figure 2*). And in many instances, only the 75 aa-long binders captured the experimentally-validated PPIs (*Table 2*). However, the 100 and 150 aa-long binder matches were enriched in GO terms that were more specific to the defense/immune response against fungi than the 75 aa-long binders (*Figure 4*). The distribution of the structural similarity scores between the binder lengths isare also very similar in most cases, yet it does appear that the 75 aa-long binders produced a disproportionately larger number of acceptable matches (bit score ≥ 50) than the other binder lengths. Interestingly, there was no clear pattern for how the binders’ length influenced the number of presumptive interacting host proteins for each individual binder (*Figure 5*).

**Figure 4.**
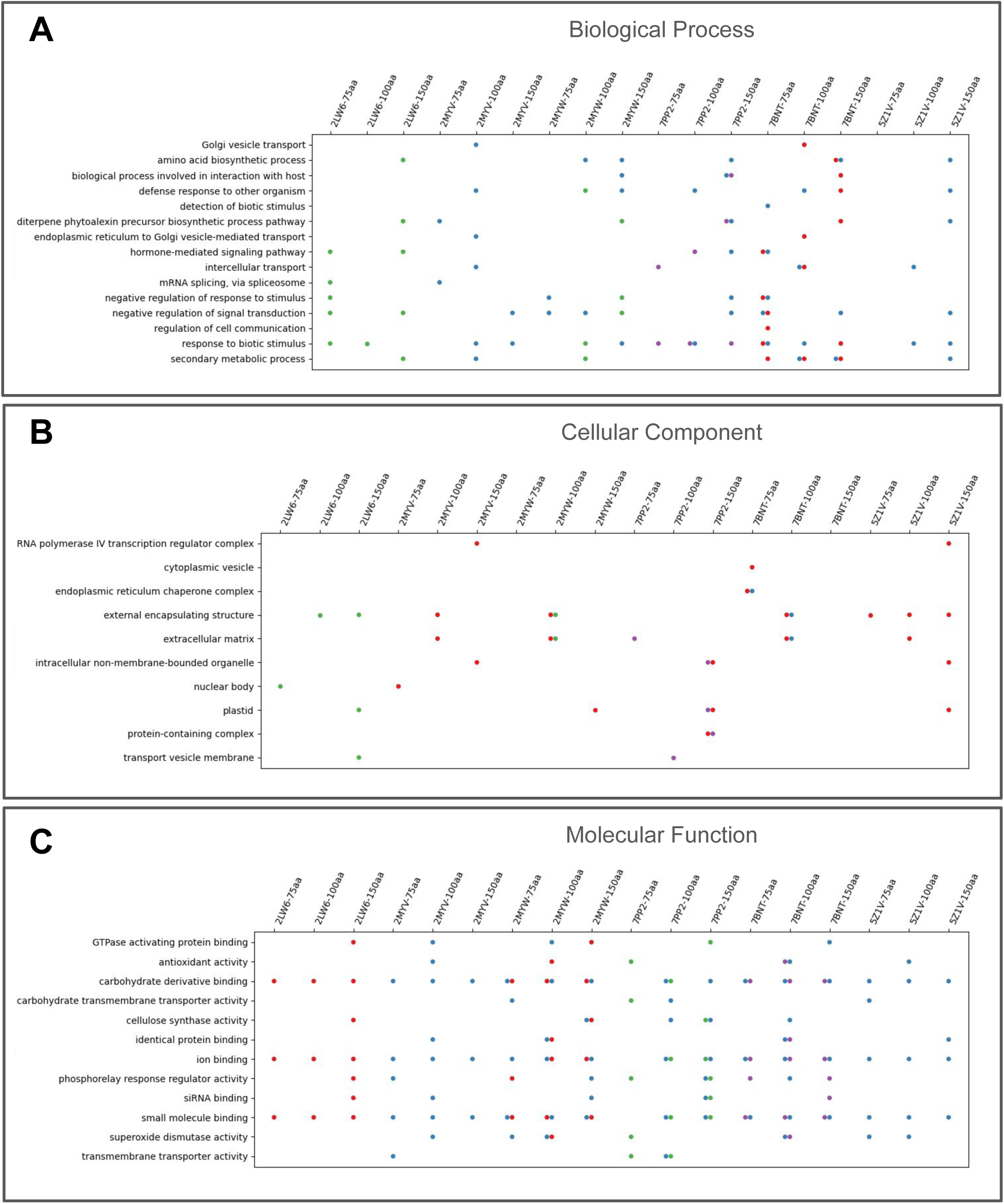
A subset of the enriched gene ontology terms for the rice proteins with structural similarity to the generated binders. The enrichments are shown by effector protein, binder length, and by the site type of the hotspots passed during diffusion: general protein binding site (blue), substrate/effector recognition site (red), active site (green), and cofactor binding site (yellow). Enrichments were performed with the Bonferroni adjustment for multiple comparisons and significance at P ≤ 0.05. The effectors are referred to by their PDB: 2LW6 (Avr-Piz-t), 2MYV (Avr1-CO39), 2MYW (Avr-Pia), 7PP2 (Avr-Pii), 7BNT (Avr-PikD), and 5Z1V (Avr-Pib).

**Figure 5.**
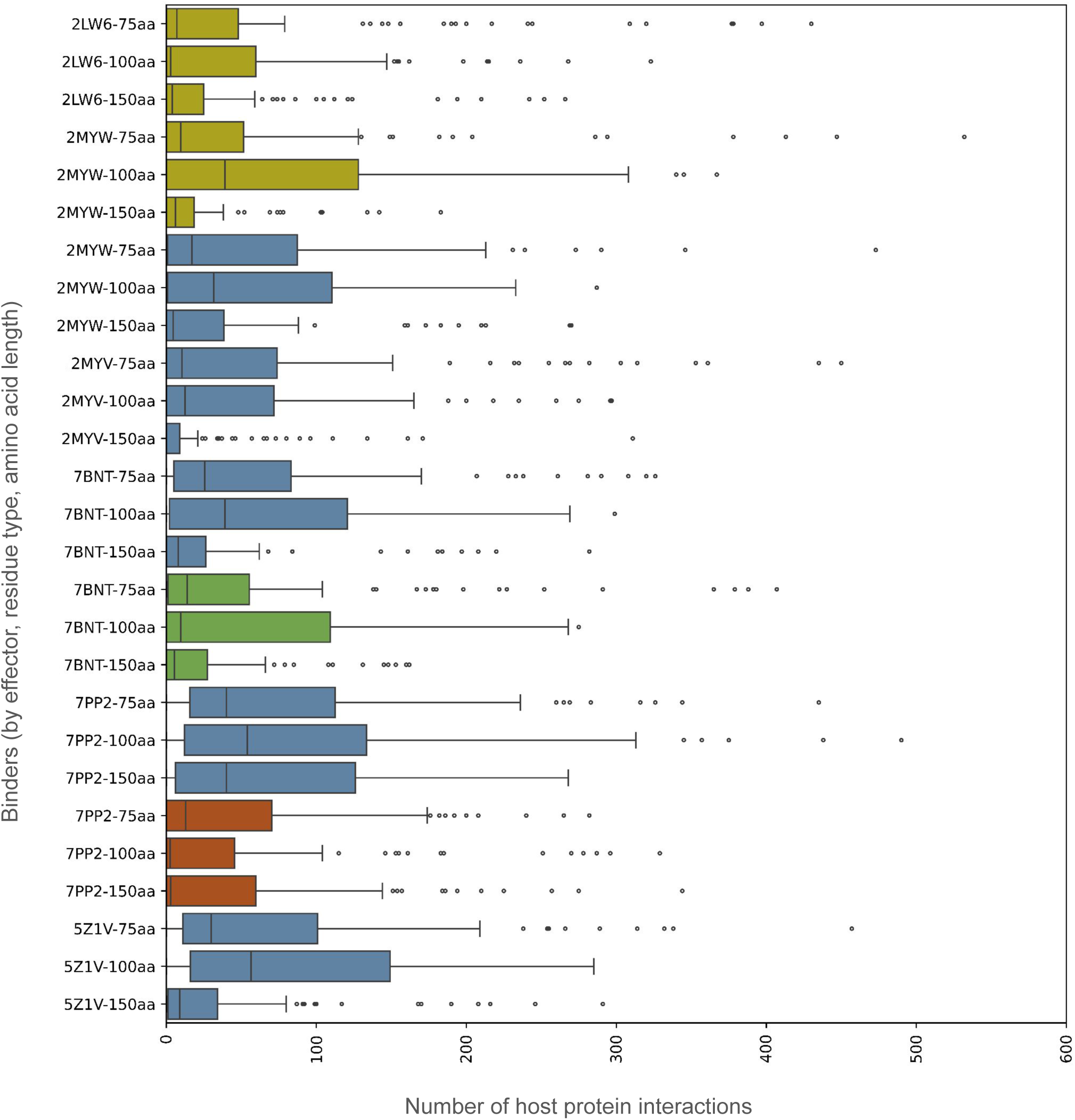
The distribution of the number of host proteins captured by each effector based on the site type passed during diffusion, and the length of the generated binder: 75, 100, or 150 amino acid (aa) residues. The site types are color-coded: blue – protein binding site, yellow – substrate recognition site, green – active/catalytic site, red – cofactor binding site. The effectors are referred to by their PDB: 2LW6 (Avr-Piz-t), 2MYV (Avr1-CO39), 2MYW (Avr-Pia), 7PP2 (Avr-Pii), 7BNT (Avr-PikD), and 5Z1V (Avr-Pib).

This finding supports the hypothesis that shorter binders introduce noise, as there are a larger number of acceptable matches with the 75 aa binders, but their functions seem to be less specific to the defense/immune response than the 100/150 aa binders’ matches. The shorter binders also appear to support the capture of more structurally diverse proteins, since the number of unique protein matches per binder was similar across the different binder lengths despite the shorter binders producing an overall larger number of protein matches. Decreasing the bit score threshold to allow for more flexibility during the structural similarity search may increase the recall for the longer binders, but could also increase the false positive rate. Generating more binders could also be an option as the authors of RFdiffusion recommend diffusing 10,000 binders for a single protein target. However, this is likely not feasible for many research groups with limited computing resources. Ultimately, the results of this validation study indicate that further evidence is needed to identify promising candidates for *in vitro* and *in vivo* validation. While the binders would capture multiple expected PPI targets for some effectors, the presumptive host protein matches for others varied by the mode of action of the input residues. For example, there was not a single binder for Avr-Pia that produced highly confident matches for all three anticipated PPIs (RGA4, RGA5, Pik1/Pik5-NP). Using protein-binding residues as the RFdiffusion input, one of the 75-residue binders aligned with a region of Pik5-NP that is structurally similar to the Heavy Metal Binding domain of the Pik-1 protein; this region facilitates the effector’s recognition in blast resistant cultivars (Guo et al., 2018). However, for the RGA4/RGA5 heterodimer, a 75-residue binder generated from protein-binding residues aligned with the apoptotic protease-activating factor domain (IPR042197) of RGA4, and the leucine rich repeat (LRR; IPR032675) domain of RGA5; only RGA5 was recovered from the binders with input recognition residues. This is significant as it corroborates with the literature that RGA4 is responsible for activating the apoptotic response pathway whereas RGA5 is only sufficient for recognition (Césari et al., 2014).

### Identifying promising candidates for *in vitro* validation

This validation study revealed two potential methods for selecting the most promising candidates. The most evident method is to take advantage of the GO term annotations and identify candidates with defense-related annotations that are over-represented in the study items. For instance, the PPI pairs could be explored on the basis of a cellular compartment of interest, or by the biological processes the effector disrupts. Of note, comparing the GO enrichments based on the active site type did reveal slight differences in the functions of the captured proteins (*Figure 4*), though with the small sample size of the experimentally validated PPIs, how to best use each active site type to further steer PPI predictions is not immediately clear. The disadvantage of this GO-based method is that it limits the exploration of unannotated or poorly annotated proteins.

As an alternative method, promising targets can be identified based on how well the binder overlaps with critical domains of the target protein. For instance, there were six 75 aa-long binders for Avr-PikD which showed structural similarity to Pik5-NP. Its homolog in rice blast resistant cultivars, Pik1, has a Heavy Metal (HMA) Binding domain which facilitates the effector’s recognition (Guo et al., 2018). In a global alignment, there is moderate structural similarity and low sequence similarity between Pik-1 and Pik5-NP (alignment length–890 residues, sequence identity–0.502, TM-score–0.6663). However, the region of Pik5-NP with structural similarity to the query binder had considerable structural similarity to the HMA domain of the Pik-1(*Figure 6*)(alignment length–67 residues, sequence identity–0.299, TM-score–0.8037). This finding provides evidence that generating these truncated protein segments can uncover important sub-structures of proteins that contribute to resistance, even if there is low sequence similarity. Such domain-based evidence could also substantiate target proteins as potential candidates for further experimentation. Other examples for query-target pairs aligned within a domain important to the host protein’s function are shown in Figure 7.

**Figure 6.**
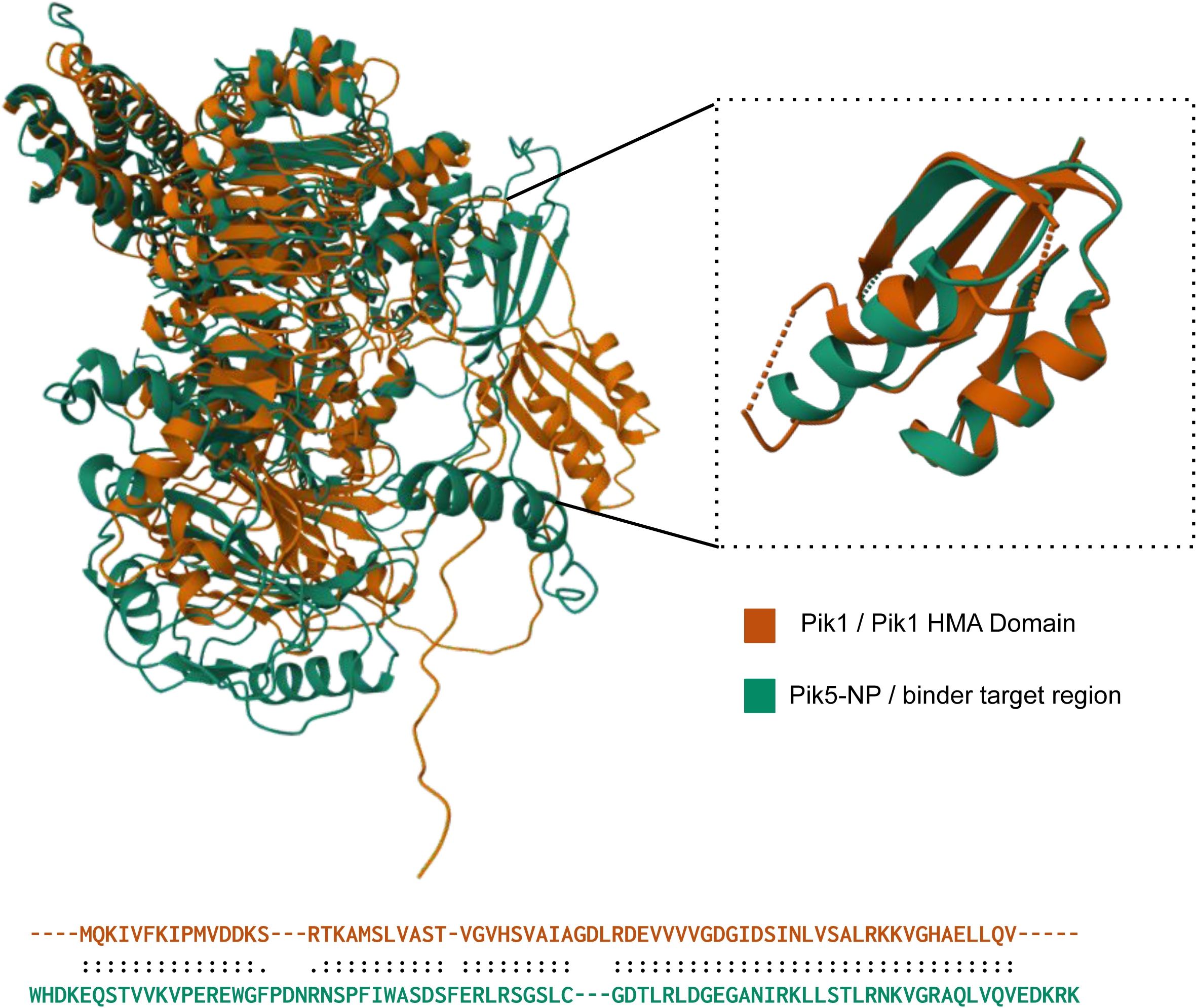
The structural alignment of the Heavy Metal Associated (HMA) domain of Pik1 (teal) with the region of Pik5-NP (orange) with structural similarity to the binders generated with RFdiffusion, using the TM-align server. A global structural alignment of the Pik1 and Pik5-NP protein indicates there is moderate structural similarity and low sequence similarity between the proteins (Alignment length = 890 residues; sequence identity = 0.502; TM-score = 0.6663). When, a structural alignment was performed using only the region that overlapped with the binder, it was found that the Pik5-NP substructure was highly similar to the HMA domain of Pik1 (Aligned length = 67 residues; RMSD = 1.81 Å; TM-score 0.80366) but had low sequence similarity (sequence identity = 0.299). Pik1 was used as the reference protein to compute the TM-score. “:” denotes aligned residue pairs of d < 5.0 Å, “.” denotes other aligned residues.

**Figure 7.**
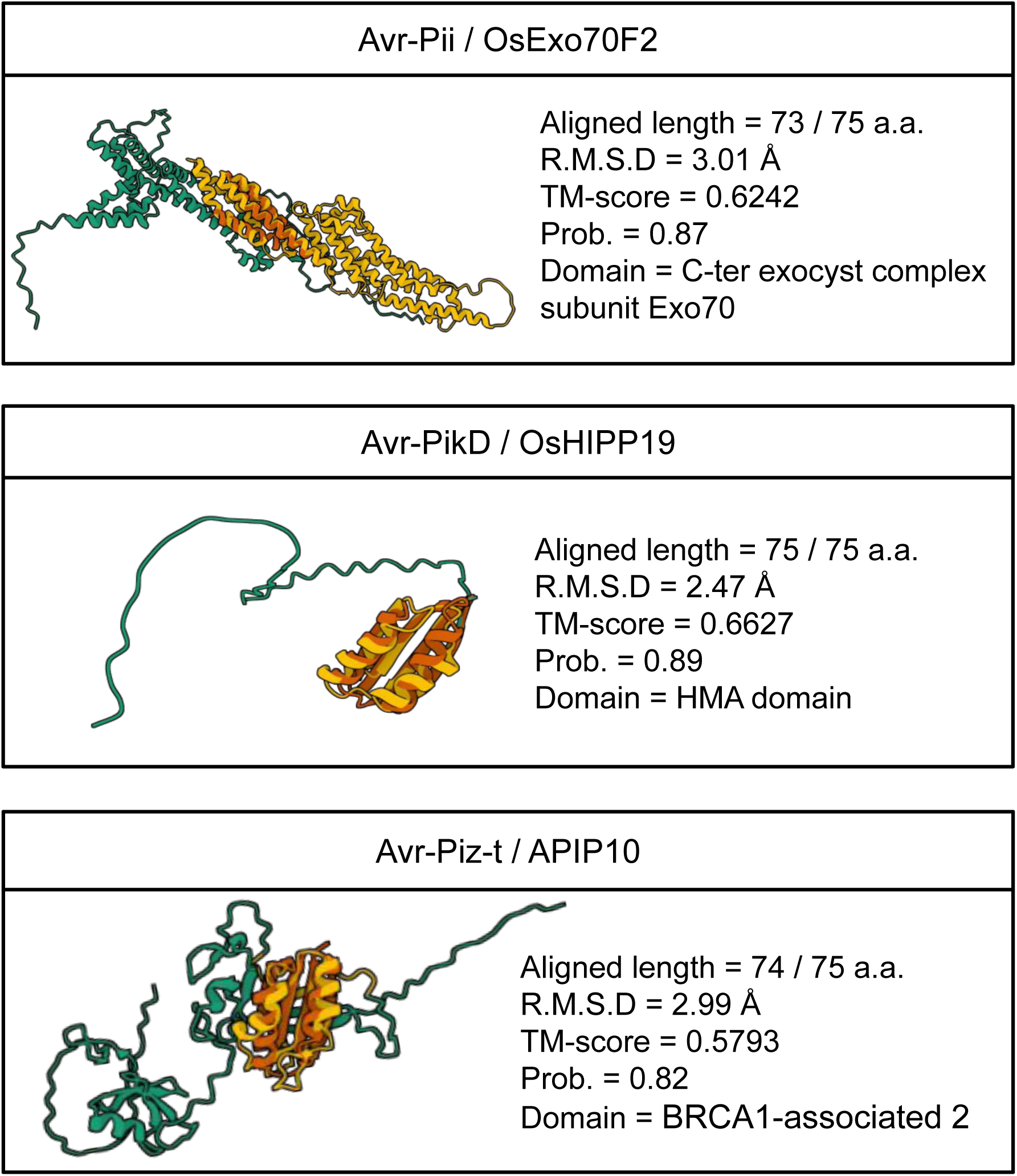
A selection of the binders and important domains of the structurally similar host proteins. For each host protein (teal), the protein domain (yellow) that contains the segment with structural similarity to the binder (orange) is shown. During the structural similarity search, the minimum coverage of the query binder was 90% (67 a.a.), and TM scores demonstrate the relatively high structural similarity of the query-target pair. FoldSeek also reports a probability (Prob.) score which reflects the probability that the matches are structurally homologous.

## Discussion

Diffusion models like RFdiffusion are gaining traction in the field of protein design for biomedical applications (Harris et al., 2024; Liu et al., 2024, Vázquez Torres et al., 2024). To the authors’ knowledge, this is the first study to exploit the generative diffusion capabilities for investigating plant-microbe interactions. The presented workflow produced a >78% recall of experimental PPIs, and adds to the growing literature on effective practices to use diffusion to study PPIs, particularly for under-represented organisms in protein structural databases. Domain knowledge is still critical to decipher PPIs, especially since this workflow could produce improbable presumptive host proteins (i.e., due to different cellular localization). Yet further exploration of the diverse structures produced during diffusion could be used to develop future research directions in plant-microbe interactions, and develop technologies to mitigate fungal infection and mycotoxin production.

This workflow could also shed new light on understudied research avenues in plant-microbe interactions. For example, more information is needed to understand how fungal pathogens employ structural homologs to ‘hijack’ the plant stress response for nutrient production, promote cell death, and other processes. (Xu et al., 2022; Shi et al., 2016). Similarly, there is limited information on how effectors are trafficked throughout the cell or degraded *in planta* (Yuen et al., 2023), or how splice variants in plants contribute to the biotic stress response against fungi (Tognacca et al., 2023). Producing *in silico* PPI predictions could help guide the research process and reduce the need for costly experimentation.

It is important to note that the binders were not validated using ProteinMPNN and AF2, as in the original publication. This step was bypassed, as preliminary trials (data not shown) found that recovering a binder that fit all the recommended quality metrics (pLDDT, pAE, pTM) was extremely difficult. This difficulty was attributed to the low representation of plant proteins in the PDB (<4%), which was used to train AF2. As per its GitHub page, the default RFdiffusion model has a bias towards producing alpha-helical structures (hence the reason for using the beta model for a better balance of structures). This imbalance is likely because extensive beta sheets are not as prominent in species like *Homo sapiens* (which encompass ∼31% of the structures available in the PDB). The workflow in this validation study instead proposes using the number of interaction hotspots between the binder and query effector, as well as the structural bit score to assess the binder structures. Ideally, more plant protein structures will be resolved in the future to improve the balance of training data in structural biology. Resolving more fungal protein structures would also be beneficial, as this study had a small sample size of fungal effectors due to the limited number of experimentally-validated PPIs wherein the effectors’ PDB structure and interaction residues were recorded in the literature.

Comparing the results of this validation study to previous studies is difficult due to differing pathogen-host systems and methodologies. For example, although the workflow captured ≥78% of the intended target proteins, the true positive rate is difficult to determine without experimental validation of the interactions between the effectors’ and the other captured proteins. Two recent studies aimed at developing structure-based methods for PPI predictions between rice and rice blast fungus - Ma et al. (2019) and Zheng et al. (2021) - tackle reporting their results in different ways. Ma et al. (2019) used a 10-fold crossover test to validate the performance of their support vector machine model at an accuracy of 90.43% for the rice-*Magnaporthe oryzae* pathosystem, though it is worth noting that this model was not assessed at a proteome-wide scale. Zheng et al. (2021) combined structural and sequence-based (domain, interlog) methods to predict PPIs between rice and *Magnaporthe oryzae* at a proteome-wide scale, and developed a scoring method to assess the potential PPIs. While the authors report a true positive rate of 20.77% (false positive rate 0.10%), the method only detected two resistance genes (pi-d2, LOC_Os06g29810; pi-ta, LOC_Os12g18360) and did not recover PPIs for most of the effectors in this study (the exception being Avr-PikD).

Also, regarding the false positive rate, it is interesting to note that the workflow did not capture host proteins that interacted indirectly (i.e., mediated through another protein). For example, Pik-1/Pik-2 form homo- and heterodimers, and both proteins are required to elicit a cell death response upon recognition of an avirulence protein (Maqbool et al., 2015). While the interactions involving Pik-1 (Pik5-NP) were recovered, interactions between Pik-2 (Pik6-NP) were not. It is not likely that this is due to substantial differences in global sequence or structural similarity between the homologs. Using the TM-align algorithm, Pik-2 and Pik6-NP appear to have higher structural and sequence similarity (alignment length-984 residues, sequence identity-0.802, TM-score-0.9351) than Pik-1 and Pik5-NP (alignment length-890 residues, sequence identity-0.502, TM-score-0.6663). These findings are in line with the literature that reports direct interactions between multiple avirulence proteins and Pik-1 (Avr-Pia, Césari et al., 2013; Avr-PikD, Kanzaki et al., 2012), whereas direct interactions between avirulence proteins and Pik2 are not reported. Kanzaki et al. (2012) even provided experimental evidence that Pik-2 and AVR-PikD do not interact directly.

Regarding other limitations, this methodology will likely find little success in identifying the gamut of non-proteinaceous resistance mechanisms in plants such as lipids (Cavaco et al., 2021) or secondary metabolites (Savignac et al., 2023). Moreover, the effectors used for validation in this study are well-characterized. Their mode of action, binding sites, and structure are well-documented in the literature (Table 2). Applying this workflow to plant pathogens that are under-represented in the literature may be more difficult, though various strategies could be used to determine protein binding sites computationally (i.e., protein language models; Schrieber et al., 2023) or through conserved domains (i.e., InterPro; Paysan-Lafosse et al., 2022). Computationally determined protein structures (i.e., from AF2 or ESMFold) could have been used instead of experimental structures from PDB. However, a thorough study on how the quality of computational protein structures affects the diffusion process is needed.

## Conclusion

As the global burden of fungal disease on crops is expected to increase due to climate change, there is a need for strategies to prevent, detect, and mitigate fungal pathogens. Leveraging the roles and interactions between proteins, which are involved in many aspects of fungal invasion and plant defense against fungi, could be indispensable to addressing this need. However, there is insufficient validated evidence for PPIs between important fungal species and most plant hosts, making a validation set very challenging. This study introduced a workflow for *in silico* predictions of fungal effectors and plant proteins guided by structural homology, and including quality metrics for assessing the outputs at nearly every stage. In all, the findings demonstrate the ability of RFdiffusion to generate structurally diverse binders that still reflect that of actual binding pairs. This research will be used to build an interaction database for major crop plant-fungal pathogen systems (i.e., *Magnaporthe-*rice, *Fusarium*-wheat, *Fusarium*-maize, etc.), providing PPIs with experimental evidence from different sources. With experimental validation forthcoming, the predictions from this protocol will be made publicly available at public databases like MaizeGDB (https://maizegdb.org/; Woodhouse et al., 2021), GrainGenes (https://wheat.pw.usda.gov/; Yao et al., 2022), and the Fusarium Protein Toolkit (https://fusarium.maizegdb.org; Kim et al., 2024). These efforts will promote research in plant-host interactions, and enable users to easily query proteins of interest, investigate their interaction PPIs, and provide visual 3D structure networks of the target protein.

## ACKNOWLEDGEMENTS

The work for this project was performed on the Atlas and Ceres high-performance clusters as part of the USDA-ARS SCINet initiative. We would like to thank the SCINet administrative staff and the Virtual Research Support Core team. This work was supported by the U.S. Department of Agriculture, Agricultural Research Service. Mention of trade names or commercial products in this publication is solely for the purpose of providing specific information and does not imply recommendation or endorsement by the U.S. Department of Agriculture. USDA is an equal opportunity provider and employer.

